# Clustered brachiopod Hox genes are not expressed collinearly and are associated with lophotrochozoan novelties

**DOI:** 10.1101/058669

**Authors:** Sabrina M. Schiemann, José M. Martín-Durán, Aina Børve, Bruno C. Vellutini, Yale J. Passamaneck, Andreas Hejnol

**Affiliations:** Sars International Centre for Marine Molecular Biology, University of Bergen, Bergen 5006, Norway; Kewalo Marine Laboratory, Pacific Biosciences Research Center, University of Hawaii, Honolulu, HI, USA

**Author notes:** Corresponding author: Andreas Hejnol.

## Abstract

Temporal collinearity is often regarded as the force preserving Hox clusters in vertebrate genomes. Studies that combine genomic and gene expression data in invertebrates would allow generalizing this observation across all animals, but are scarce, particularly within Lophotrochozoa (e.g., snails and segmented worms). Here, we use two brachiopod species –*Terebratalia transversa*, *Novocrania anomala*– to characterize the complement, cluster and expression of their Hox genes. *T. transversa* has an ordered, split cluster with ten genes (*lab*, *pb*, *Hox3*, *dfd*, *scr*, *lox5*, *antp*, *lox4*, *post2*, *post1*), while *N. anomala* has nine (missing *post1*). Our *in situ* hybridization, qPCR and stage specific transcriptomic analyses show that brachiopod Hox genes are neither strictly temporally nor spatially collinear; only *pb* (in *T. transversa*), *Hox3* and *dfd* (in both brachiopods) show staggered mesodermal expression. The spatial expression of the Hox genes in both brachiopod species correlates with their morphology and demonstrates cooption of Hox genes in the chaetae and shell fields, two major lophotrochozoan morphological novelties. The shared and specific expression of a subset of Hox genes, *Arx* and *Zic* orthologs in chaetae and shell-fields between brachiopods, mollusks, and annelids supports the deep conservation of the molecular basis forming these lophotrochozoan hallmarks. Our findings challenge that collinearity alone preserves lophotrochozoan Hox clusters, indicating that additional genomic traits need to be considered in understanding Hox evolution.

## Introduction

Hox genes are transcription factors that bind to regulatory regions via a helix-turn-helix domain to enhance or suppress gene transcription [1, 2]. Hox genes were initially described in the fruit fly *Drosophila melanogaster* [3, 4] and later on in vertebrates [5–7] and the nematode *Caenorhabditis elegans* [8]. In all these organisms, Hox genes were shown to provide a spatial coordinate system for cells along the anterior-posterior axis [9]. Remarkably, the Hox genes of these organisms are clustered in their genomes and exhibit a staggered spatial [3] and temporal [10, 11] expression during embryogenesis that corresponds to their genomic arrangement [3, 12, 13]. These features were used to classify Hox genes in four major orthologous groups –anterior, Hox3, central and posterior Hox genes– and were proposed to be ancestral attributes to all bilaterally symmetrical animals [1, 13, 14].

However, the study of the genomic arrangements and expression patterns of Hox genes in a broader phylogenetic context has revealed multiple deviations from that evolutionary scenario. Hox genes are prone to gains [15–17] and losses [18–21], and their arrangement in a cluster can be interrupted, or even completely disintegrated [22–25]. Furthermore, the collinear character of the Hox gene expression can fade temporally [24, 26, 27] and/or spatially [28]. Hox genes have also diversified their roles during development, extending beyond providing spatial information [29]. In many bilaterian embryos, Hox genes are expressed during early development, well before the primary body axis is patterned [26, 30–32]. They are also involved in patterning different tissues [33] and have been often recruited for the evolution and development of novel morphological traits, such as vertebrate limbs [34, 35], cephalopod funnels and arms [28], and beetle horns [36].

It is thus not surprising that Hox genes show diverse arrangements regarding their genomic organization and expression profiles in the Spiralia [37], a major animal clade that includes a high disparity of developmental strategies and body organizations [38–42]. A striking example is the bdelloid rotifer *Adineta vaga,* which belongs to the Gnathifera, the possible sister group to all remaining Spiralia [41, 42]. As a result of their reduced tetraploidy, its Hox complement includes 24 genes, albeit it lacks posterior Hox genes and a Hox cluster [43]. The freshwater flatworms *Macrostomum lignano* and *Schmidtea mediterranea* also lack a Hox cluster [44, 45] and parasitic flatworms have undergone extensive Hox gene losses, likely associated with their particular life style [21]. Interestingly, the limpet mollusk *Lottia gigantea* [16] shows a well-organized Hox cluster. Other mollusks (e.g. the pacific oyster *Crassostrea gigas*) and the segmented annelid *Capitella teleta* exhibit organized split Hox clusters [46, 47]. On the other hand, the cephalopod mollusk *Octopus bimaculiodes* has lost several Hox genes and lacks a Hox cluster [22]; and the clitellate annelids *Helobdella robusta* and *Eisenia fetida* do not show a Hox cluster and have greatly expanded some of the Hox classes [16, 17].

Although Hox gene expression is known for a handful of spiralian species [26, 44, 46, 48–58], the relationship between genomic organization and expression domains is known for only three of them, namely the annelids *C. teleta* and *H. robusta*, and the planarian *S. mediterranea*. Consistent with the lack of a Hox cluster, *H. robusta* and *S. mediterranea* show neither temporal nor spatial collinearity [44, 54–56]. Conversely, *C. teleta*, which has an organized, broken cluster, does exhibit these features [46]. In general, these observations suggest that the presence of collinearity – in particular, temporal collinearity– could be associated with the retention of a more or less intact spiralian Hox cluster, as it seems the case for the vertebrate cluster [14, 23, 59, 60]. However, more studies combining genomic and expression information, and including the vast spiralian morphological diversity, are essential to draw robust conclusions about Hox gene evolution and regulation in Spiralia and Metazoa [61]. These studies would also allow to test if hypotheses about the correlation between collinearity and cluster organization as observed in deuterostomes [23] stand true for protostomes.

Here, we present a comprehensive study of the genomic arrangement and expression of Hox genes in Brachiopoda, a lineage of the Spiralia whose origins date back to the Lower Cambrian [62]. Brachiopods are marine, sessile, filter-feeding animals. They are protected by two dorsoventral mineralized shells and reproduce by external fertilization, often developing through an intermediate, free-living larval stage [63]. In this study, we use two brachiopod species –the ‘articulate’ *Terebratalia transversa* and the ‘inarticulate’ *Novocrania anomala*– that respectively belong to the two major brachiopod lineages, thus allowing the reconstruction of putative ancestral characters for Brachiopoda as a whole (Figure 1A). By transcriptomic and genomic sequencing we demonstrate that the Hox complement consists of ten Hox genes in *T. transversa* and nine in *N. anomala*. In addition, the ten *Hox* genes of *T. transversa* are ordered in a split Hox cluster that differs from the genomic arrangement reported for the brachiopod *Lingula anatina* [personal communication, Luo and 64]. We show that Hox gene expression is restricted to the ‘trunk’ region of the larva, and is overall neither temporally nor spatially collinear. However, the genes *pb* (only in *T. transversa*), *Hox3* and *dfd* show spatially collinear expression in the mesoderm of both brachiopod species. Additionally, the Hox genes *lab*, *scr*, *antp* and *post1* appear to be associated with the development of two brachiopod features: the chaetae and the shell-forming epithelium. Altogether, our findings demonstrate that the presence of a split Hox cluster in the Brachiopoda is likely not associated with a temporally collinear expression of Hox genes, which differs from the hypothesized correlation between temporal collinearity and the retention of the vertebrate Hox cluster [14, 23, 59, 60] and suggests that alternative/additional genomic forces might shape Hox clusters during spiralian evolution, such as low genomic rearrangement frequency.

**Figure 1.**
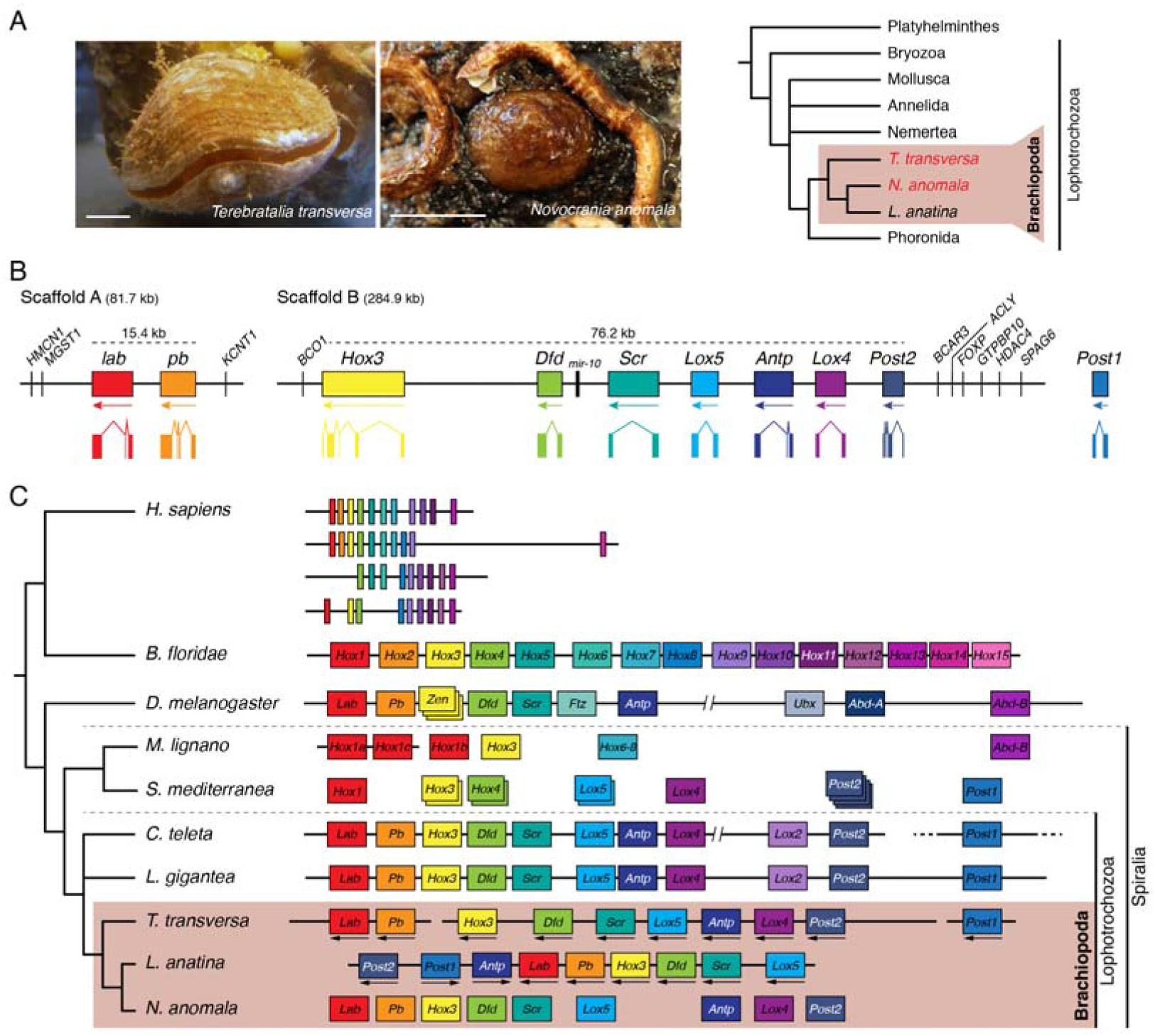
Genomic organization of Hox genes in Brachiopoda. (**A**) Images of an adult *T. transversa* and *N. anomala*, and phylogenetic position of these species within Brachiopoda and Lophotrochozoa. (**B**) The ten Hox genes of *T. transversa* are ordered along three genomic scaffolds and are flanked by external genes (vertical lines; gene orthology is based on best blast hit). Thus, *T. transversa* has a split Hox cluster composed of three sub-clusters. No predicted ORFs were identified between the Hox genes in scaffold A and B. A colored box represents each Hox gene, and below each box there is the direction of transcription and the exon-intron composition. The genomic regions containing Hox genes are represented in scale. (**C**) The genomic organization of brachiopod Hox genes in a phylogenetic context (adapted from [22]). The genomic order of Hox genes in *T. transversa* is similar to that observed in other spiralians (e.g. *Capitella teleta* and *Lottia gigantea*), which suggests that the translocation of the posterior Hox cluster upstream to lab is a lineage-specific feature of *L. anatina* (in *T. transversa* and *L. anatina* the arrows below the genes show the direction of transcription; conformation of the Hox cluster in L. anatina was kindly provided by N. Satoh and Y.-J. Luo). The low contiguity of the draft genome assembly of *N. anomala* hampered recovering genomic linkages between the identified Hox genes. Each ortholog group is represented by a particular color.

## Results

### *The* Hox *gene complement of* T. transversa *and* N. anomala

Transcriptomic and genomic searches resulted in the identification of ten Hox genes in *T. transversa*. In the brachiopod *N. anomala*, we identified seven Hox genes in the transcriptome and two additional fragments corresponding to a Hox homeodomain in the draft genome assembly. Attempts to amplify and extend these two genomic sequences in the embryonic and larval transcriptome of *N. anomala* failed, suggesting that these two Hox genes might be expressed only during metamorphosis and/or in the adult brachiopod. Maximum likelihood orthology analyses resolved the identity of the retrieved Hox genes (Figure supplementary 1). The ten Hox genes of *T. transversa* were orthologous to *labial* (*lab*), *proboscipedia* (*pb*), *Hox3*, *deformed* (*dfd*), *sex combs reduced* (*scr*), *lox5*, *antennapedia* (*antp*), *lox4*, *post2* and *post1*. The nine Hox genes identified in *N. anomala* corresponded to *lab*, *pb*, *Hox3*, *dfd*, *scr*, *lox5*, *antp*, *lox4*, and *post2*.

### *Genomic organization of* Hox *genes in* T. transversa *and* N. anomala

We used the draft assemblies of *T. transversa* and *N. anomala* genomes to investigate the genomic arrangement of their Hox genes. In *T. transversa*, we identified three scaffolds containing Hox genes (Figure 1B). Scaffold A spanned 81.7 kb and contained *lab* and *pb* in a genomic region of 15.4 kb, flanked by other genes with no known linkage to the Hox cluster in other animals. Scaffold B was the longest (284.8 kb) and included *Hox3*, *dfd*, *scr*, *lox5*, *antp*, *lox4* and *post2*, in this order (Figure 1B) including the micro RNA *mir-10* between *dfd* and *scr*. As in scaffold A, other genes flanked the Hox genes, which occupied a genomic region of 76.2 kb. Finally, *post1* aligned to various short scaffolds. We could not recover any genomic linkage between the identified Hox genes in *N. anomala* due to the low contiguity (N50 of 3.5 kb) of the draft genome assembly. Altogether, these data demonstrate that *T. transversa* has a split Hox cluster broken into three sub-clusters, each of them with an organized arrangement. Importantly, the potential genomic disposition of these three sub-clusters is similar to that observed in other spiralians, such as *C. teleta* and *L. gigantea* (Figure 1C), which suggests that the lineage leading to the brachiopod *L. anatina* experienced genomic rearrangements that modified the ordered and linkage of the Hox genes.

### Hox *gene expression in* T. transversa

To investigate the presence of temporal and/or spatial collinearity in the expression of the clustered Hox genes in *T. transversa*, we first performed whole-mount *in situ* hybridizations in embryos from blastula to late, competent larval stages (Figure 2).

**Figure 2.**
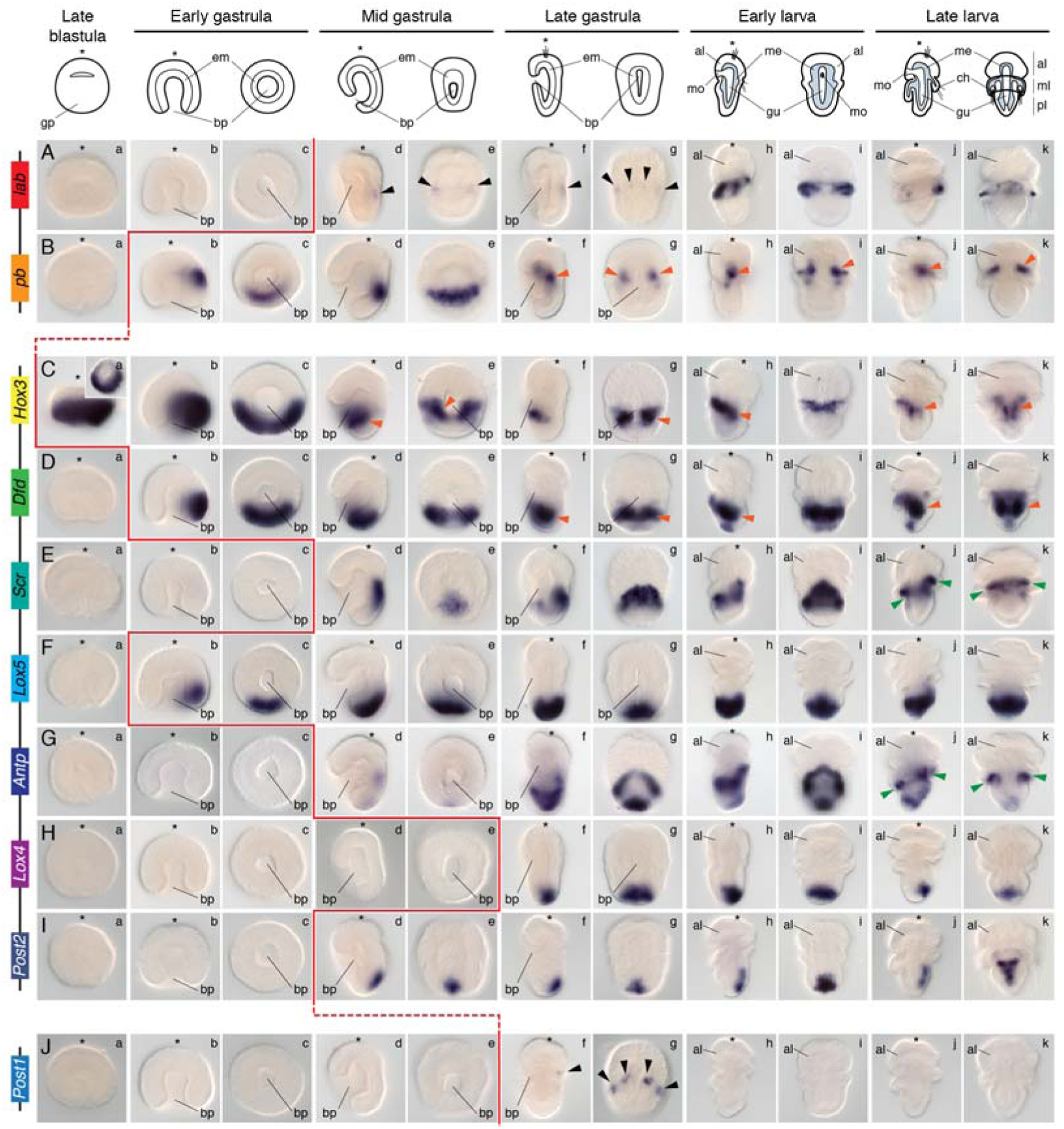
Expression of Hox genes in *T. transversa*. (**A**–**J**) Whole mount *in situ* hybridization of each Hox gene during embryonic and larval stages in *T. transversa*. The Hox genes *lab* and *post1* are expressed during chaetae formation. The genes *pb*, *Hox3* and *dfd* are collinearly expressed along the mantle and pedicle mesoderm. The Hox genes *scr* and *antp* are expressed in the periostracum, the shell-forming epithelium. *Lox5*, *Lox4* and *post2* are expressed in the posterior ectoderm of the pedicle lobe. See main text for a detailed description of each expression pattern. Black arrowheads indicate expression in the chaetae sacs. Orange arrowheads highlight mesodermal expression. Green arrowheads indicate expression in the periostracum. The genomic organization of the Hox genes is shown on the left. On top, schematic representations of each analyzed developmental stage on its respective perspective. In these schemes, the blue area represents the mesoderm. Drawings are not to scale. The red line indicates the onset of expression of each Hox gene based on *in situ* hybridization data. The blastula stage is a lateral view (inset is a vegetal view). The other stages are in a lateral view (left column) and dorsoventral view (right column). The asterisk demarcates the animal/anterior pole. al, apical lobe; bp, blastopore; ch, chaetae; em, endomesoderm; gp, gastral plate; gu, gut; me, mesoderm; ml, mantle lobe; mo, mouth; pl, pedicle lobe.

#### Anterior Hox genes

The anterior Hox gene *lab* was first detected in the mid gastrula stage in two faint bilaterally symmetrical dorsal ectodermal domains (Figure 2Ad, Ae). In late gastrulae, *lab* expression consisted of four dorsal ectodermal clusters that corresponded to the position where the chaetae sacs form (Figure 2Af, Ag). In early larva, the expression was strong and broad in the mantle lobe (Figure 2Ah, Ai), and in late larvae it became restricted to a few mantle cells adjacent to the chaetae sacs (Figure 2Ij, Ik). These cells do not co-localize with tropomyosin, which labels the muscular mesoderm of the larva (Figure 3A). This suggests that *lab* expressing cells are likely ectodermal, although we cannot exclude localization in non-muscular mesodermal derivates.

**Figure 3.**
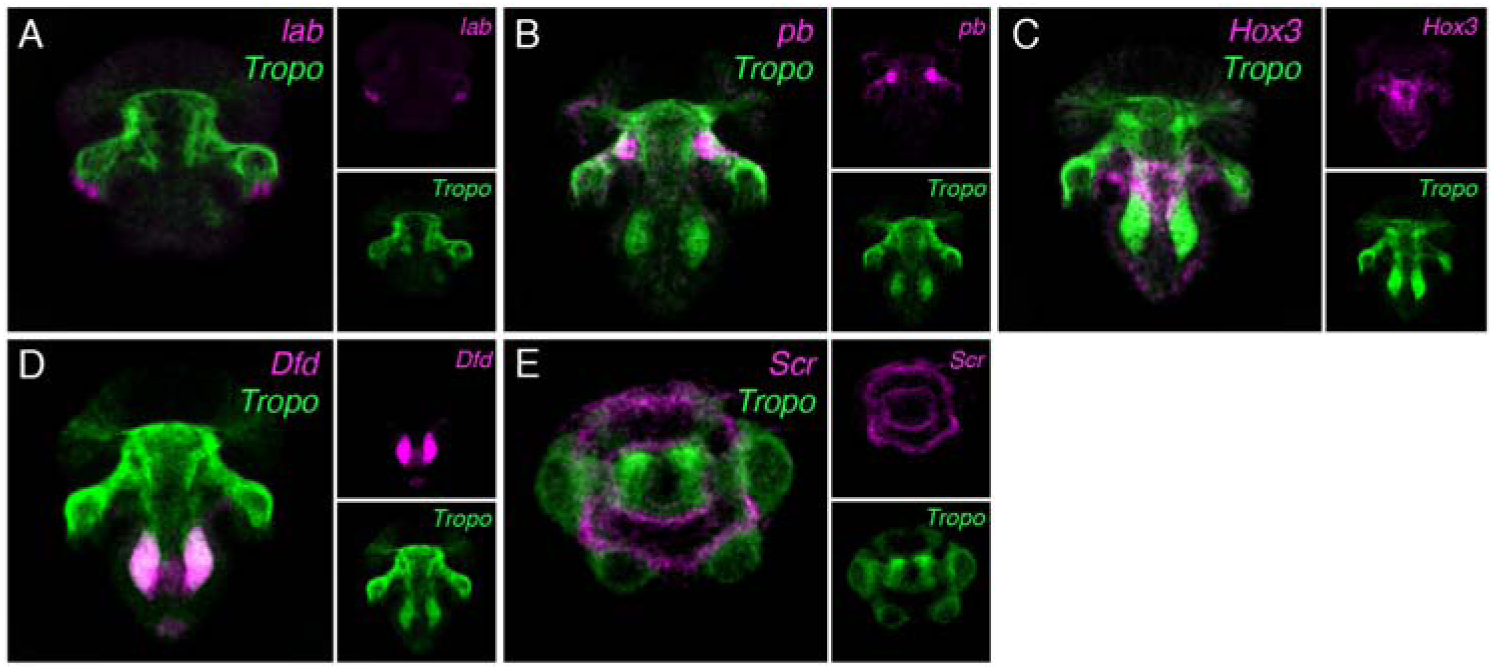
Hox expression in mesoderm and periostracum of *T. transversa*. (**A**–**E**) Double fluorescent *in situ* hybridization of *lab*, *pb*, *Hox3*, *dfd* and *scr* with tropomyosin (Tropo, in green) in late larval stages of *T. transversa*. (**A**) The gene *lab* is expressed in relation to the chaetae sacs, but does not overlap with the tropomyosin-expressing mesoderm. (**B**–**D**) The Hox genes *pb*, *Hox3* and *Dfd* show spatial collinearity along the mantle and pedicle mesoderm. (**E**) The gene *scr* is expressed in the periostracum, which is the epithelium that forms the shell.

The Hox gene *pb* was first detected asymmetrically on one lateral of the ectoderm of the early gastrula (Figure 2Bb, Bc). In the mid gastrula, the ectodermal domain located dorsally and extended as a transversal stripe (Figure 2Bd, Be). Remarkably, this domain disappeared in late gastrula embryos, where *pb* was detected in the anterior mantle mesoderm (Figure 2Bf, Bg). This expression was kept in the early and late larva (Figure 2Bh–Bk; Figure 3B)

#### Hox3

The gene *Hox3* was detected already in blastula embryos in a circle of asymmetric intensity around the gastral plate (Figure 2Ca). In early gastrulae, Hox3 is restricted to one half of the vegetal one, which is the prospective posterior side (Figure 2Cb, Cc). With axial elongation, *Hox3* becomes expressed in the anterior mantle mesoderm and in the ventral ectoderm limiting the apical and mantle lobe (Figure 2Cd, Ce). This expression is maintained in late gastrula stages and in the early larva (Figure 2Cf–Ci). In the late larva, *Hox3* is detected in part of ventral, internal mantle ectoderm and in the most anterior part of the pedicle mesoderm (Figure 2Cj, Ck; Figure 3C)

#### Central Hox genes

The Hox gene *dfd* was asymmetrically expressed on one side of the vegetal pole of the early gastrula of *T. transversa* (Figure 2Db, Dc). This expression was maintained in the mid gastrula, and corresponded to the most posterior region of the embryo (Figure 2Dd, De). In the late gastrula, *dfd* becomes strongly expressed in the posterior mesoderm (Figure 2Df, Dg). In the early larva, the expression remained in the pedicle mesoderm, but new domains in the posterior ectoderm and in the anterior, ventral pedicle ectoderm appear (Figure 2Dh, Di). These expression domains are also observed in the late larva (Figure 2Dj, Dk; Figure 3D).

The central Hox gene *scr* was first expressed in the medial dorsal ectoderm of the mid gastrula (Figure 2Ed, Ee). In late gastrula stages, the expression expanded towards the ventral side, forming a ring (Figure 2Ef, Eg). In the early larva, *scr* was detected in a ring encircling the most anterior ectoderm of the pedicle lobe and extending anteriorly on its dorsal side (Figure 2Eh, Ei). With the outgrowth of the mantle lobe in the late larva, the expression became restricted to the periostracum, the internal ectoderm of the mantle lobe that forms the shell (Figure 2Ej, Ek; Figure 3E).

The Hox gene *Lox5* is expressed on one side of the early gastrula (Figure 2Fb, Fc). During axial elongation, the expression became restricted to the most posterior ectoderm of the embryo (Figure 2Fd–Fg). This domain remained constant in larval stages, where it was expressed in the whole posterior ectoderm of the pedicle lobe (Figure 2Fh–Fk).

The *antp* gene is weakly detected at the mid gastrula stage, in one posterior ectodermal domain and one dorsal ectodermal patch (Figure 2Gd, Ge). In the late gastrula, the posterior expression is maintained and the dorsal domain extends ventrally, encircling the embryo (Figure 2Gf, Gg). These two domains remained in the larvae: the ectodermal anterior-most, ring-like domain localized to the periostracum, and the posterior domain limited to the most posterior tip of the larva (Figure 2Gh–Gk).

The Hox gene *Lox4* is first detected in the dorsal, posterior end of the late gastrula and early larva (Figure 2Hf–Hi). In the late larva, *Lox4* is expressed dorsally and posteriorly, although it is absent from the most posterior end (Figure 2Hj, Hk).

#### Posterior Hox genes

The posterior Hox gene *post2* was first detected in mid gastrula stages at the posterior tip of the embryo (Figure 2Id, Ie). This expression was maintained in late gastrulae (Figure 2If, Ig). In early larva, *post2* expression extended anteriorly and occupied the dorso-posterior midline of the pedicle lobe (Figure 2Ih, Ii). In late, competent larvae, *post2* was detected in a T-domain in the dorsal side of the pedicle ectoderm (Figure 2Ij, Ik).

The Hox gene *post1* was transiently detected in late gastrula stages in the four mesodermal chaetae sacs (Figure 2Jf, Jg).

We verified the absence of temporal collinearity in the expression of the Hox genes in *T. transversa* by quantitative real-time PCR and comparative stage-specific RNA-seq data (Figure supplementary 2).

### Hox *gene expression in* N. anomala

In order to infer potential ancestral Hox expression domains for the Brachiopoda, we investigated the expression of the nine Hox genes of *N. anomala* during embryogenesis and larval stages (Figure 4).

**Figure 4.**
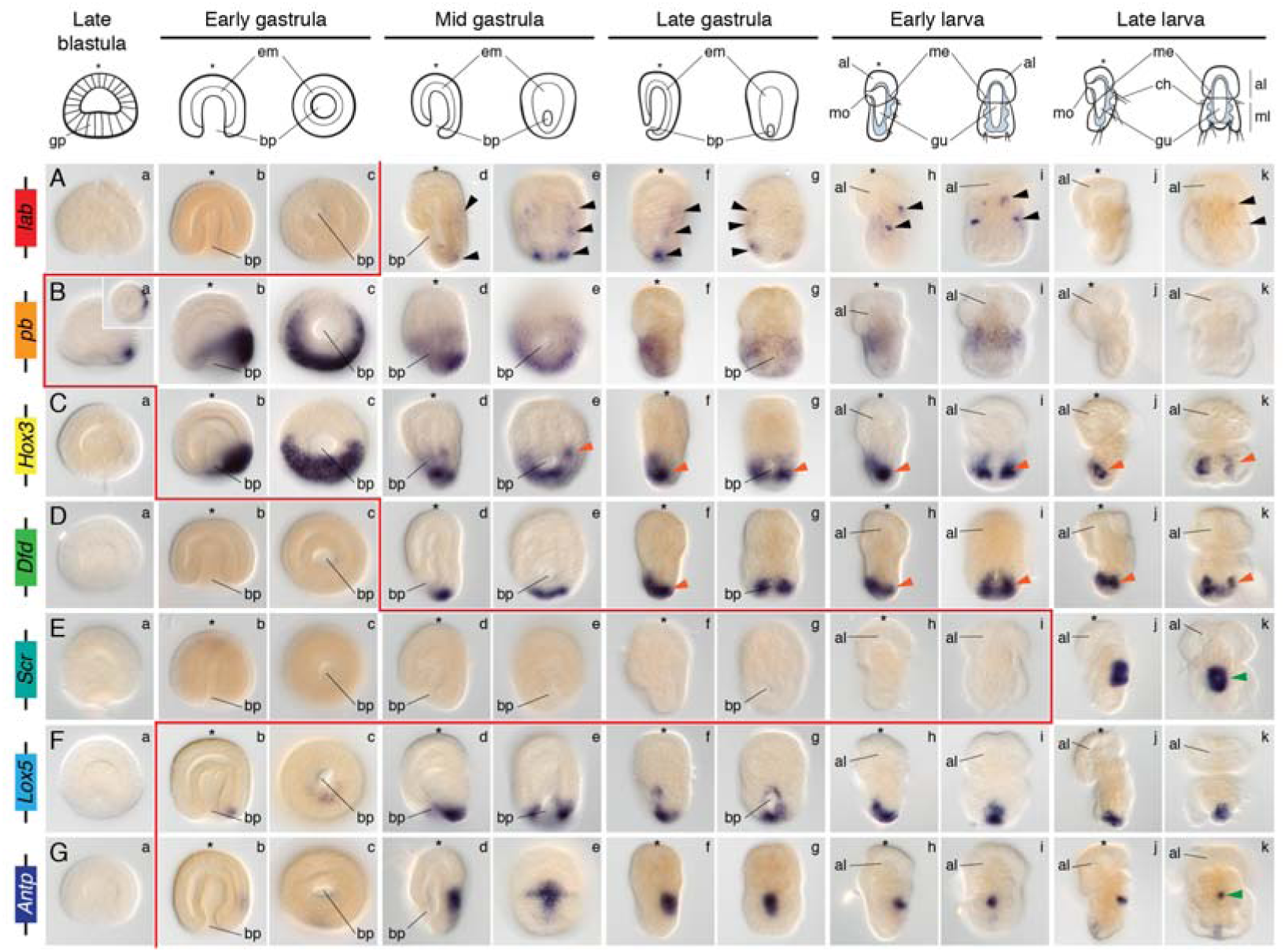
Expression of Hox genes in *N. anomala*. (**A**–**G**) Whole mount *in situ* hybridization of the Hox genes during embryonic and larval stages in *N. anomala*. The gene *lab* is expressed in the chaetae. The Hox genes *Hox3* and *dfd* are collinearly expressed in the mantle mesoderm. The genes *scr* and *antp* are expressed in the prospective shell-forming epithelium. The genes *pb* and *Lox5* are detected in the ectoderm of the mantle lobe. The genes *Lox4* and *post2* were not detected in transcriptomes and cDNA during embryonic stages. See main text for a detailed description of each expression pattern. Black arrowheads indicate expression in the chaetae sacs. Orange arrowheads highlight mesodermal expression. Green arrowheads indicate expression in the periostracum. On top, schematic representations of each analyzed developmental stage on its respective perspective. In these schemes, the blue area represents the mesoderm. Drawings are not to scale. The red line indicates the onset of expression of each Hox gene based on *in situ* hybridization data. The blastula stage is a lateral view (inset is a vegetal view). The other stages are in a lateral view (left column) and dorsoventral view (right column). The asterisk demarcates the animal/anterior pole. al, apical lobe; bp, blastopore; ch, chaetae; em, endomesoderm; gp, gastral plate; gu, gut; me, mesoderm; ml, mantle lobe; mo, mouth.

#### Anterior Hox genes

The Hox gene *lab* was first detected at the mid gastrula stage in three bilaterally symmetrical ectodermal cell clusters that appear to correlate with the presumptive site of chaetae sac formation (Figure 4Ad, Ae). The expression in the most posterior pair was stronger than in the two most anterior ones. This expression was maintained in the late gastrula (Figure 4Af, Ag). In larval stages, *lab* was detected in the two most anterior chaetae sacs of the mantle lobe (Figure 4Ah, Ai), expression that fainted in late larvae (Figure 4Aj, Ak).

The Hox gene *pb* was asymmetrically expressed already at blastula stages, in the region that putatively will rise to the most posterior body regions (Figure 4Ba). With the onset of gastrulation, the expression of *pb* extended around the vegetal pole, almost encircling the whole blastoporal rim (Figure 4Bb, Bc). During axial elongation, *pb* was first broadly expressed in the region that forms the mantle lobe (Figure 4Bd, Be) and later on the ventral mantle ectoderm of the late gastrula (Figure 4Bf, Bg). In early larvae, *pb* was detected in the anterior ventral mantle ectoderm (Figure 4Bh, Bi). This domain was not detected in late, competent larvae (Figure 4Bj, Bk).

#### Hox3

The Hox gene *Hox3* was asymmetrically detected around half of the vegetal pole of the early gastrulae (Figure 4Cb, Cc). In mid gastrulae, the expression almost encircled the whole posterior area and the blastoporal rim (Figure 4Cd). In addition, a domain in the mid-posterior mesoderm became evident (Figure 4Ce). By the end of the axial elongation, *Hox3* was strongly expressed in the posterior mesoderm and weakly in the ventral posterior mantle ectoderm (Figure 4Cf, Cg). Noticeably, the posterior most ectoderm did not show expression of *Hox3*. This expression pattern was maintained in early and late larval stages (Figure 4Ch–Ck).

#### Central Hox genes

The central Hox gene *dfd* was first detected in the posterior ectodermal tip of mid gastrulae (Figure 4Dd, De). In late gastrula stages, *dfd* was expressed in the posterior ectodermal end (Figure 4Df) and in the posterior mesoderm (Figure 4Dg). Early larvae showed expression of *dfd* in the posterior mesoderm and posterior mantle ectoderm (Figure 4Dh, Di). This expression remained in late larvae, although the most posterior ectodermal end was devoid of expression (Figure 4Dj, Dk).

The Hox gene *scr* was only detected in late larval stages, in a strong dorsal ectodermal domain (Figure 4Ej, Ek).

The gene *Lox5* was detected asymmetrically around half of the blastoporal rim in early gastrula stages (Figure 4Fb, Fc). During axial elongation, the expression progressively expanded around the blastoporal rim (Figure 4Fd, Fe) and limited to the ventral midline (Figure 4Ff, Fg). In the larvae, *Lox5* was expressed in the ventral, posterior-most midline (Figure 4Fh–Fk).

The Hox gene *antp* was first expressed asymmetrically in one lateral side of the early gastrula (Figure 4Gj, Gk). In the mid gastrula, *antp* was detected in the dorsal ectodermal mantle in a cross configuration: dorsal midline and the mantle cells closer to the apical-mantle lobe boundary (Figure 4Gd, Ge). In late gastrulae, *antp* was only expressed in a mid-dorsal ectodermal region (Figure 4Gf, Gg). This expression pattern was also observed in early larval stages, although the size of the domain reduced (Figure 4Gh, Gi). In late larvae, antp was detected in a small mid-dorsal patch and a weak ventro-posterior ectodermal domain (Figure 4Gj, Gk).

We could neither identify nor amplify *Lox4* in a transcriptome and cDNA obtained from mixed embryonic and larval stages, suggesting that either it is very transiently and weakly expressed during embryogenesis or it is only expressed in later stages (metamorphosis and adulthood).

#### Posterior Hox genes

The only posterior Hox gene present in *N. anomala*, *post2*, could not be amplified in cDNA obtained from mixed embryonic and larval stages, suggesting that it is not expressed –or at least expressed at really low levels– during these stages of the life cycle. The absence of larval expression of *Lox4* and *post2* could be related to the lack of the pedicle lobe of craniiform brachiopod larva, which is a characteristics of the lineage [65, 66].

## Discussion

### The brachiopod Hox complement and the evolution of Hox genes in Spiralia

Our findings on *T. transversa* and *N. anomala* reveal an ancestral brachiopod Hox gene complement consistent with what has been hypothesized to be ancestral for Spiralia and Lophotrochozoa on the basis of degenerate PCR surveys [15, 67–69]. This ancient complement comprises eight Hox genes – *lab*, *pb*, *Hox3*, *Dfd*, *Scr*, *Lox5*, *Lox4* and *Post2* – and has been confirmed by genomic sequencing of representative annelids and mollusks [16, 22, 47], rotifers and platyhelminthes [21, 43–45] and the linguliform brachiopod *L. anatina* [64]. While *T. transversa* and *L. anatina* (N. Satoh and Y.-J. Luo, personal communication) have retained this ancestral Hox complement, *N. anomala* has apparently lost *Post1* (Figure 1).

Our genomic information shows that the Hox cluster of *T. transversa* is split in three parts, with *lab* and *pb* separate from the major cluster and *Post1* also on a separate scaffold (Figure 1B). Overall, the cluster extends over 100 kb, which is significantly shorter than those of other lophotrochozoans, such as *C. teleta* (~345kb) [46] and *L. gigantea* (~455 kb) [16]. Its compact size is related to short intergenic regions and introns, comparable to the situation observed in vertebrate Hox clusters [23]. The order and orientation of the Hox genes in *T. transversa* is preserved and more organized than in the Hox cluster reported for the brachiopod *L. anatina*, which exhibits a genomic rearrangement that placed a portion of the cluster upstream *lab* and in reverse orientation [64]. Indeed, the split Hox clusters reported so far in lophotrochozoan taxa exhibit all different conformations, indicating that lineage-specific genomic events have shaped Hox gene clusters in Spiralia.

### Non-collinearity of Hox expression in T. transversa despite the presence of a split cluster

The analysis of Hox clustering in different animal species together with the temporal and spatial expression patterns of their Hox genes grounded the hypotheses that the regulatory elements required for their collinearity –mostly temporal– maintain the clustered organization of the vertebrate Hox genes and possibly other animals [13, 23, 59–61, 70, 71] (Figure supplementary 3). Although there are cases in which spatial collinearity is displayed in the absence of a cluster, as in the appendicularian chordate *O. dioica* [24], all investigated clustered Hox genes show at least one type of collinearity that could account for their genomic organization [23, 61] (Figure 6). Since there are exceptions to the spatial collinearity in vertebrates, for instance *Hoxa2* and *Hoxb2* are expressed more anteriorly than *Hox1* genes in the vertebrate hindbrain [72], temporal collinearity is seen as a manifestation of Hox clustering. But whether temporal collinearity is the agent keeping the cluster together, e.g. through enhancer sharing [73], is still subject of debate.

Within Spiralia, this evolutionary scenario appears to be supported by the staggered temporal and spatial expression of the Hox genes in the split cluster of the annelid *C. teleta* [46]. In the other investigated spiralians, there is only either genomic information (e.g. the mollusks *L. gigantea* and *C. gigas*) or expression analysis (e.g. the mollusks *G. varia*, *Haliotis asinina*) [16, 47, 52, 58]. Most of these gene expression studies have demonstrated coordinated spatial or temporal expression of Hox genes along the anteroposterior axis of the animal [48, 49, 74] or in organ systems, such the nervous system [52, 58]. However, the absence of studies that can reveal a correlation between the expression of Hox genes and their genomic organization in these animals hampers the reconstruction of the putative mechanisms that preserve Hox clusters in Lophotrochozoa, and thus prevent generalizations about possible scenarios of Hox cluster evolution across all animals.

Our findings robustly demonstrate that the Hox genes of the split Hox cluster of *T. transversa* overall show neither strictly spatial nor temporal collinearity (Figures 2, 3), and lack quantitative collinearity [61], as it has been shown for example in mouse [75]. These observations are also supported by the absence of a coordinated spatial and temporal expression of the Hox genes in *N. anomala* (Figure 4). Although a general trend of spatial collinearity is present (e.g. the posterior Hox genes are expressed in posterior tissues), the early expression of *Hox3* breaks temporal collinearity in *T. transversa*, while it is *pb* that becomes first expressed in *N. anomala*. In both species, the gene *Lox5* is also expressed before *Scr*, as it is also the case in the annelid *N. virens* [74]. Ectodermal spatial collinearity is absent in the two brachiopods even when considering the future location of the larval tissues after metamorphosis [76, 77]. The most anterior class gene *lab* is exclusively expressed in the chaetae of *T. transversa* and *N. anomala*, and thus is not affiliated with anterior neural or foregut tissues as in other lophotrochozoans, such as annelids [46, 78]. Similarly, the most posterior Hox gene, *Post1*, is very transiently expressed in the chaetae sacs, which occupy a mid-position in the larval body. We only detected a strict spatial collinearity in the staggered expression of the Hox genes *pb*, *Hox3* and *Dfd* along the anterior-posterior axis of the developing larval mesoderm in both *T. transversa* and *N. anomala* (Figure 5).

**Figure 5.**
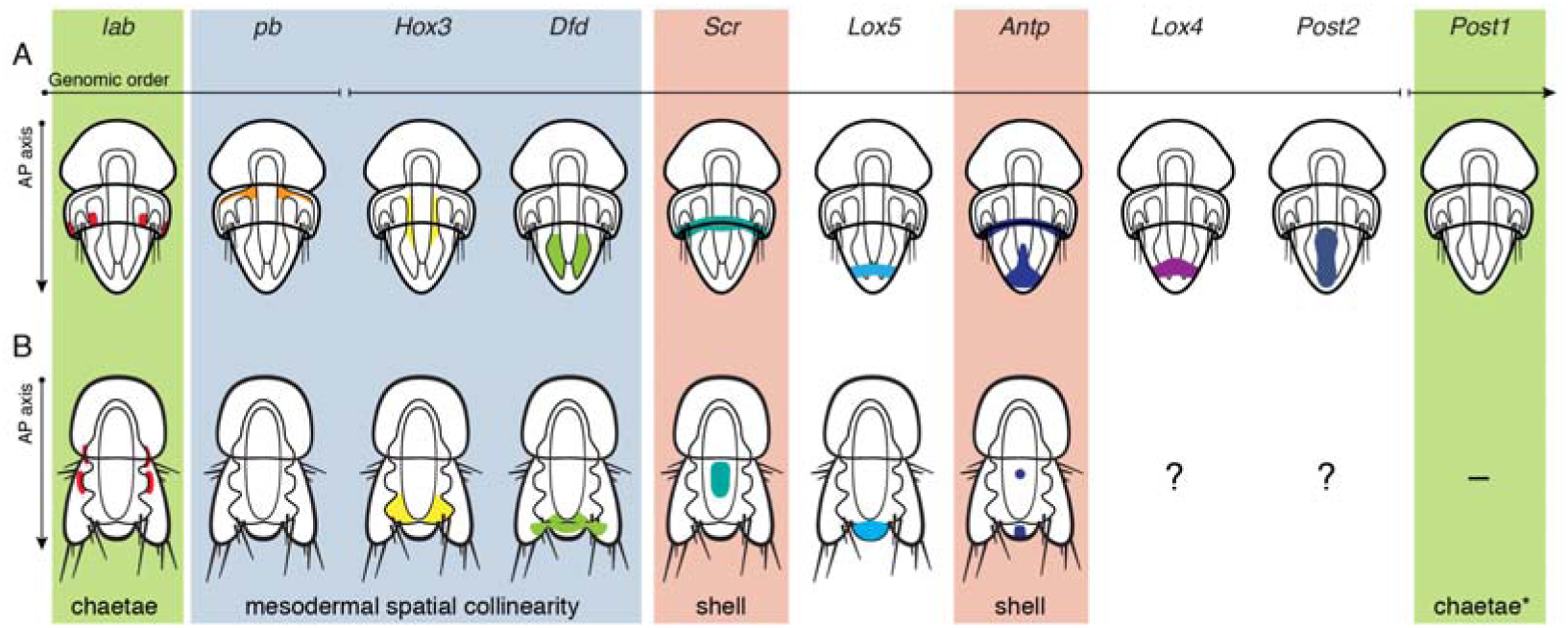
Summary of Hox gene expression in *T. transversa* and *N. anomala*. (**A**, **B**) Schematic drawings of late larvae of *T. transversa* and *N. anomala* depicting the expression of each Hox gene. The Hox genes *pb* (not in *N. anomala*), *Hox3* and *dfd* show staggered expression, at least in one of their domains, associated with the mesoderm (light blue box). In both brachiopods, the genes *scr* and *antp* are expressed in the periostracum, or the shell-forming epithelium (red boxes) and lab and post1 are associated to the developing chaetae (green boxes; asterisk in *post1*: *post1* is expressed in the chaetae only during late embryonic stages, not in the mature larva, and only in *T. transversa*). The expression of *Lox4* and *post2* in *N. anomala* could not be determined in this study. The gene *post1* is missing in *N. anomala*. Drawings are not to scale.

Altogether, the absence of a strict temporal and spatial collinearity in the brachiopod *T. transversa*, albeit the presence of a split Hox cluster, indicates that temporal collinearity is likely not the underlying factor keeping spiralian Hox genes clustered, as it seems to be the case in vertebrates [14, 23, 59–61]. Therefore, alternative mechanisms might need to be considered. In this regard, why do Hox clusters split in different positions between related species, as seen for instance in brachiopods (this study) and drosophilids [79], but still display similar expression profiles? This might indicate that the control of expression in large split Hox clusters relies more on gene-specific short-range transcriptional control than on a global, coordinated regulation, as seen in the small Hox vertebrate clusters [23, 75, 80]. The conservation of Hox clusters in many animals could then be a consequence of the general conservation of syntenic relationships in their genomes (Figure 6). Our findings thus highlight the necessity of further detailed structure-function analyses of spiralian Hox clusters to better understand the intricate evolution of the genomic organization and regulation of Hox genes in metazoans.

**Figure 6.**
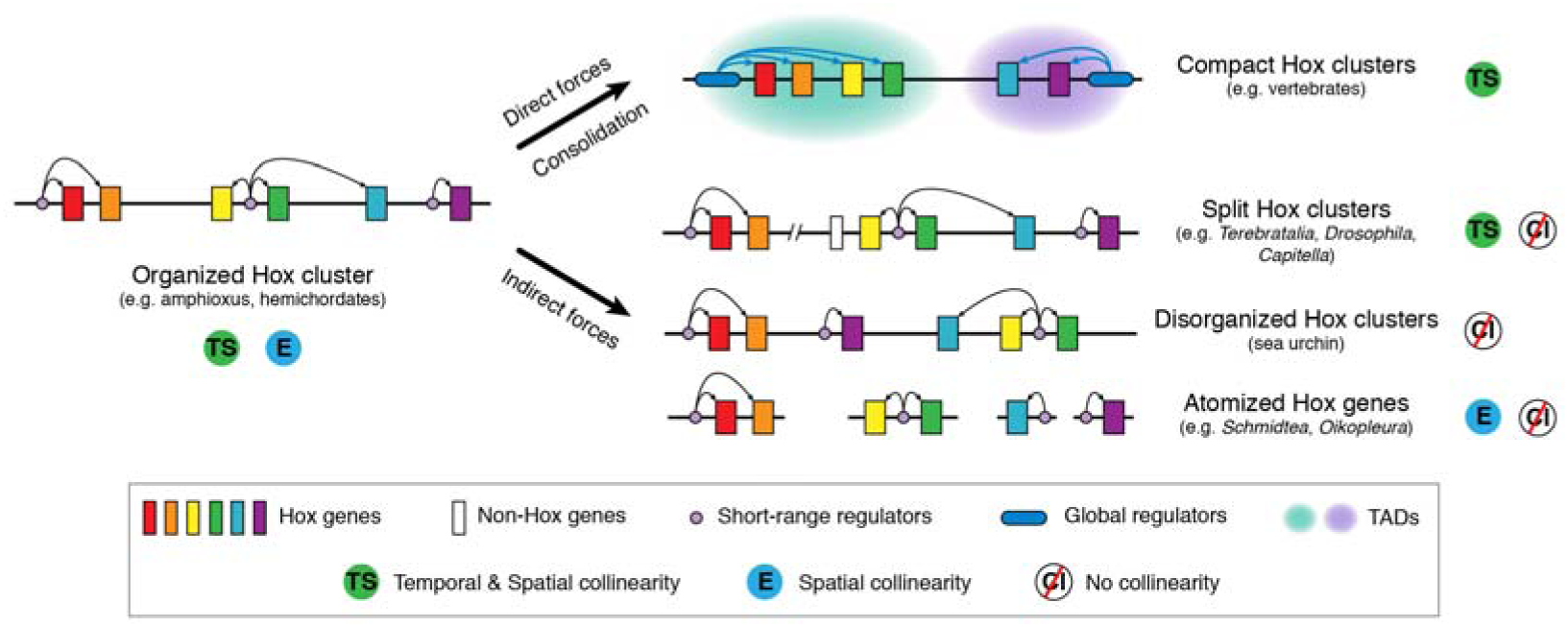
Hox gene cluster evolution. The absence of temporal collinearity in the ordered, split Hox cluster of the brachiopod *T. transversa* weakens the scenario that temporal collinearity is a major force keeping Hox genes together. Alternatively, we propose that the ancestral Hox cluster was organized and regulated by short-range regulators, which elaborated a collinear expression, as observed in amphioxus (temporal and spatial collinearity) and hemichordates (only spatial collinearity). Direct selective pressures could have promoted genomic changes that led to the evolution of compact, tightly regulated, collinear Hox clusters (e.g. in vertebrates). In other animal lineages, the original organized Hox cluster would become broken, rearranged or atomized indirectly, as a result of general genome-wide events. In these cases, the conservation/loss of the ancestral regulatory elements, together with the evolution of new ones associated with novel morphologies, would have influence the collinear expression of Hox genes. Under this scenario, collinearity is only one of multiple direct or indirect factors affecting Hox clustering.

### Recruitment of Hox genes for patterning lophotrochozoan chaetae and shell fields

The bristle-like chaetae (or setae) of annelids and brachiopods, and shell valves in mollusks and brachiopods are the most prominent hard tissues found in lophotrochozoan spiralians [63] and provide fossilized hallmarks of the Cambrian explosion [81]. It has been already recognized that the ultrastructural morphology of the brachiopod and annelid chaetae is nearly identical [82–84] and with the placement of brachiopods as close relatives of annelids and mollusks [85], the homology of these structures appeared more likely [86]. In this context, the anterior Hox gene *lab* is expressed in the chaetae of *Chaetopterus* sp. [26] and *Post1* is expressed in the chaetae of *C. teleta*, *P. dumerilii* and *N. virens* [46, 74]. Our results show that similarly, *lab* and *Post1* are expressed specifically in the chaetal sacs of the brachiopods *T. transversa* and *N. anomala* (Figures 2, 4) and follow the different arrangement of the chaetae in both species. Further evidence of a common, and probably homologous, molecular profile comes from the expression of the homeodomain gene *Aristaless-like* (*Arx*) and the zinc finger *Zic*. These genes are expressed at each chaetae sac territory in the *Platynereis* larva [87], in *Capitella teleta* [88], and also in the region of the forming chaetae sac territories in *T. transversa* (Figure supplementary 4). Therefore, the expression of the Hox genes *lab* and *Post1* and the homeodomain gene *Arx* indicate that similar molecular signature underlays the development of chaetae in annelids and brachiopods. This, together with the evident and striking morphological similarities shared by brachiopod and annelid chaetae, support considering these two structures homologous, and thus, common lophotrochozoan innovations. This would be consistent with placing the iconic Cambrian fossil *Wiwaxia*, which contains chaetae, as a stem group lophotrochozoan [89].

The protective shell is a mineralized tissue present in brachiopods and mollusks. In the gastropod mollusk *G. varia*, the Hox genes *lab*, *Post1* and *Post2* are first expressed in the shell field, and later is *Dfd* [57]. In *H. asinina* also *lab* and *Post2* are related to shell formation [52]. In brachiopods, *Dfd* is associated to the adult shell in *L. anatina* [64]. During embryogenesis of *T. transversa* and *N. anomala*, however, only *Scr* and *Antp* are expressed in the shell fields, but not *lab* or *Post1*, which are expressed in the chaetae sacs. This could support the homology of the chitin-network that is formed at the onset of brachiopod and mollusk shell fields. However, the different deployment of Hox genes in the shell fields of brachiopods and mollusks might indicate that these genes do not have an ancient role in the specification of the shell-forming epithelium. However, their consistent deployment during shell development might reflect a more general, conserved role in shaping the shell fields according to their position along the anterior posterior axis.

## Conclusions

In this study, we characterize the Hox gene complement of the brachiopods *T. transversa* and *N. anomala*, and demonstrate the last common ancestor to all brachiopods likely had ten Hox genes (*lab*, *pb*, *Hox3*, *dfd*, *scr*, *Lox5*, *antp*, *Lox4*, *post2*, *post1*). Noticeably, brachiopod Hox genes do not show global temporal and spatial collinearity, albeit *T. transversa* exhibits an ordered, split Hox cluster. Only the genes *pb* (in *T. transversa*), *Hox3* and *dfd* (in both brachiopods) show spatial collinearity in the ‘trunk’ mesoderm. In addition, the Hox genes *lab* and *post1*, as well as the homeobox *Arx*, are expressed in the developing chaetae, as also described for other annelid species [46, 53, 74]. These molecular similarities, together with evident morphological resemblances [83], support considering brachiopod and annelid chaetae homologous structures and reinforce considering the fossil *Wiwaxia* as a stem group lophotrochozoan [89]. Altogether, our findings challenge a scenario in which temporal collinearity is the major force preserving Hox clusters [12, 14, 23, 60, 61], and indicate that alternative/additional genomic mechanisms might account for the great diversity of Hox gene arrangements observed in extant animals.

## Material and Methods

### Animal cultures

Gravid adults of *Terebratalia transversa* (Sowerby, 1846) were collected around San Juan Island, Washington, USA and *Novocrania anomala* (Müller, 1776) around Bergen, Norway. Animal husbandry, fertilization and larval culture were conducted following previously published protocols [90–92].

### Hox *cluster reconstruction in* T. transversa *and* N. anomala

Male gonads of *T. transvesa* and *N. anomala* were preserved in RNAlater (Life Technologies) for further genomic DNA (gDNA) isolation. Paired end and mate pair libraries of 2 kb and 5 kb insert sizes of *T. transversa* gDNA were sequenced using an Illumina HiSeq2000 platform. First we trimmed Illumina adapters with Cutadapt 1.4.2 [93]. Then, we assembled the paired end reads into contigs, scaffolded the assembly with the mate pair reads, and closed the gaps using Platanus 1.21 [94]. The genomic scaffolds of *T. transversa* including *Hox* genes are published on GenBank with the accession numbers KX372775 and KX372776. Paired end libraries of *N. anomala* gDNA were sequenced using an Illumina HiSeq2000 platform. We removed Illumina adapters as above and assembled the paired end reads with MaSuRCA 2.2.1 [95].

### Gene isolation

Pooled samples of *T. transversa* and *N. anomala* embryos at different developmental stages (cleavage, blastula, gastrula, mid gastrula, late gastrula, early larva, and late/competent larva) were used for RNA isolation and Illumina sequencing (NCBI SRA; *T. transversa* accession SRX1307070, *N. anomala* accession SRX1343816). We trimmed adapters and low quality reads from the raw data with Trimmomatic 0.32 [96] and assembled the reads with Trinity 2.0.6 [97]. *Hox* genes were identified by BLAST searches on these transcriptomes and their respective draft genomes (see above). First-strand cDNA template (SuperScript^™^, Life Technologies) of mixed embryonic stages was used for gene-specific PCR. RACE cDNA of mixed embryonic stages was constructed with SMARTer RACE cDNA Amplification Kit (Clontech) and used to amplify gene ends when necessary. All fragments were cloned into the pGEM-T-Easy vector (Promega) and sequenced at the University of Bergen sequencing facility. *T. transversa* and *N. anomala Hox* gene sequences were uploaded to GenBank (accession numbers KX372756–KX372774).

### Orthology analyses

*Hox* gene sequences of a representative selection of bilaterian lineages (Supplementary Table S1) were aligned with MAFFT v.7 [98]. The multiple sequence alignment, which is available upon request, was trimmed to include the 60 amino acids of the homeodomain. ProtTest v.3 [99] was used to determine the best fitting evolutionary model (LG+G+I). Orthology analyses were conducted with RAxML v.8.2.6 [100] using the autoMRE option. The resulting trees were edited with FigTree and Illustrator CS6 (Adobe).

### Gene expression analyses

*T. transversa* and *N. anomala* embryos at different embryonic and larval stages were fixed in 4% paraformaldehyde in sea water for 1 h at room temperature. All larval stages were relaxed in 7.4% magnesium chloride for 10 min before fixation. Fixed samples were washed several times in phosphate buffer saline (PBS) with 0.1% tween-20 before dehydration through a graded methanol series and storage in 100% methanol at −20 °C. Single colorimetric whole mount *in situ* hybridization were carried out following an established protocol (detailed protocol available in Protocol Exchange: doi:10.1038/nprot.2008.201) [101, 102]. Double fluorescent *in situ* hybridizations were conducted as described elsewhere [103]. Representative stained specimens were imaged with bright field Nomarski optics using an Axiocam HRc connected to an Axioscope Ax10 (Zeiss). Fluorescently labeled embryos were mounted in Murray’s clearing reagent (benzyl alcohol: benzyl benzoate, 1:2) and imaged under a SP5 confocal laser-scanning microscope (Leica). Images and confocal z-stacks were processed with Fiji and Photoshop CS6 (Adobe) and figure panels assembled with Illustrator CS6 (Adobe). Contrast and brightness were always adjusted to the whole image, and not to parts of it.

### *Quantitative* Hox *gene expression in* T. transversa

Thousands of synchronous *T. transversa* embryos collected at 14 specific stages (oocytes, 8h mid blastula, 19h late blastula, 24h moving late blastula, 26h early gastrula, 37h asymmetric gastrula, 51h bilateral gastrula, 59h bilobed, 68h trilobed, 82h early larva (first chaetae visible), 98h late larva (long chaetae, eye spots), 131h competent larva, 1d juvenile, 2d juvenile) were pooled together and preserved in RNAlater (Life Technologies). Total RNA was isolated with Trizol Reagent (Life Technologies). For quantitative real time PCR, total RNA was DNAse treated and preserved at −80 °C. Gene specific primers bordering an intron splice-site and defining an amplicon of 80-150 bp sizes were designed for each gene (Supplementary Table S2). Expression levels of two technical replicates performed in two biological replicates were calculated based on absolute quantification units. For comparative stage-specific transcriptomic analyses, total RNA was used for constructing Illumina single end libraries and sequenced in four lanes of a HiSeq 2000 platform. Samples were randomized between the lanes. To estimate the abundance of transcripts per stage, we mapped the single end reads to the transcriptome of *T. transversa* with Bowtie, calculated expression levels with RSEM, and generated a matrix with TMM normalization across samples by running Trinity’s utility scripts. Expression levels obtained after quantitative real-time PCR and comparative stage-specific transcriptomics were plotted with R.

## Acknowledgements

We thank the crew of the “Centennial” boat and office stuff at Friday Harbor Laboratories (USA) and the crew of the “Hans Brattström” and “Aurelia” boats at the Espeland Marine Station (Norway) for their invaluable help during animal collections. We also thank Daniel Thiel and Anlaug Boddington for their help with animal collections and spawnings, Daniel Chourrout for his valuable comments on early versions of this manuscript, and Kevin Kocot for the access to entoproct transcriptomes. We are indebted to Yi-Jyun Luo and Nori Satoh for sharing unpublished data from the *Lingula anatina* genome assembly Version 2. The trip to Friday Harbor Laboratories was funded by a Meltzer Fond grant. The research conducted in this study was funded by the Sars Centre core budget.

## Author contributions

A.H. designed the study. A.H., S.M.S. and J.M.M.D. conducted the gene isolation and *in situ* hybridization studies. J.M.M.D. performed the gene orthology analyses. A.H., J.M.M.D., Y.P. and B.V. collected the stage-specific samples of *T. transversa* embryos. A.B. and J.M.M.D. isolated the genomic DNA of *T. transversa* and *N. anomala*. J.M.M.D. and B.V. did the draft genome assemblies and S.M.S. analyzed the Hox genomic organization. J.M.M.D. performed the stage-specific RNA isolations; A.B. did the quantitative real time PCR experiments, and B.V. conducted the analysis of the stage-specific transcriptomes. A.H. and J.M.M.D. wrote the manuscript. All authors discussed the data and edited the text.

## References

1. McGinnis W, Krumlauf R: Homeobox genes and axial patterning. Cell 1992, 68(2):283–302.

2. Pearson JC, Lemons D, McGinnis W: Modulating Hox gene functions during animal body patterning. Nat Rev Genet 2005, 6(12):893–904.

3. Lewis EB: A gene complex controlling segmentation in Drosophila. Nature 1978, 276(5688):565–570.

4. McGinnis W, Levine MS, Hafen E, Kuroiwa A, Gehring WJ: A conserved DNA sequence in homoeotic genes of the *Drosophila* Antennapedia and bithorax complexes. Nature 1984, 308(5958):428–433.

5. Carrasco AE, McGinnis W, Gehring WJ, De Robertis EM: Cloning of an *X. laevis* gene expressed during early embryogenesis coding for a peptide region homologous to Drosophila homeotic genes. Cell 1984, 37(2):409–414.

6. McGinnis W, Garber RL, Wirz J, Kuroiwa A, Gehring WJ: A homologous protein-coding sequence in drosophila homeotic genes and its conservation in other metazoans. Cell 1984, 37(2):403–408.

7. McGinnis W, Hart CP, Gehring WJ, Ruddle FH: Molecular cloning and chromosome mapping of a mouse DNA sequence homologous to homeotic genes of *Drosophila*. Cell 1984, 38(3):675–680.

8. Costa M, Weir M, Coulson A, Sulston J, Kenyon C: Posterior pattern formation in *C. elegans* involves position-specific expression of a gene containing a homeobox. Cell 1988, 55(5):747–756.

9. Akam M: Hox and HOM: homologous gene clusters in insects and vertebrates. Cell 1989, 57(3):347–349.

10. Dollé P, Izpisúa-Belmonte JC, Falkenstein H, Renucci A, Duboule D: Coordinate expression of the murine Hox-5 complex homoeobox-containing genes during limb pattern formation. Nature 1989, 342(6251):767–772.

11. Izpisúa-Belmonte J, Falkenstein H, Dollé P, Renucci A, Duboule D: Murine genes related to the Drosophila AbdB homeotic genes are sequentially expressed during development of the posterior part of the body. The EMBO journal 1991, 10(8):2279.

12. Duboule D, Morata G: Colinearity and functional hierarchy among genes of the homeotic complexes. Trends Genet 1994, 10(10):358–364.

13. Lemons D, McGinnis W: Genomic Evolution of Hox Gene Clusters. Science 2006, 313(5795):1918–1922.

14. Garcia-Fernàndez J: The genesis and evolution of homeobox gene clusters. Nat Rev Genet 2005, 6(12):881–892.

15. de Rosa R, Grenier JK, Andreeva T, Cook CE, Adoutte A, Akam M, Carroll SB, Balavoine G: Hox genes in brachiopods and priapulids and protostome evolution. Nature 1999, 399(6738):772–776.

16. Simakov O, Marletaz F, Cho SJ, Edsinger-Gonzales E, Havlak P, Hellsten U, Kuo DH, Larsson T, Lv J, Arendt D et al: Insights into bilaterian evolution from three spiralian genomes. Nature 2013, 493(7433):526–531.

17. Zwarycz AS, Nossa CW, Putnam NH, Ryan JF: Timing and Scope of Genomic Expansion within Annelida: Evidence from Homeoboxes in the Genome of the Earthworm *Eisenia fetida*. Genome Biol Evol 2016, 8(1):271–281.

18. Aboobaker A, Blaxter M: Hox gene evolution in nematodes: novelty conserved. Curr Opin Genet Dev 2003, 13(6):593–598.

19. Aboobaker AA, Blaxter ML: Hox Gene Loss during Dynamic Evolution of the Nematode Cluster. Current biology: CB 2003, 13(1):37–40.

20. Smith FW, Boothby TC, Giovannini I, Rebecchi L, Jockusch EL, Goldstein B: The Compact Body Plan of Tardigrades Evolved by the Loss of a Large Body Region. Current biology: CB 2016, 26(2):224–229.

21. Tsai IJ, Zarowiecki M, Holroyd N, Garciarrubio A, Sanchez-Flores A, Brooks KL, Tracey A, Bobes RJ, Fragoso G, Sciutto E et al: The genomes of four tapeworm species reveal adaptations to parasitism. Nature 2013, 496(7443):57–63.

22. Albertin CB, Simakov O, Mitros T, Wang ZY, Pungor JR, Edsinger-Gonzales E, Brenner S, Ragsdale CW, Rokhsar DS: The octopus genome and the evolution of cephalopod neural and morphological novelties. Nature 2015, 524(7564):220–224.

23. Duboule D: The rise and fall of Hox gene clusters. Development 2007, 134(14):2549–2560.

24. Seo HC, Edvardsen RB, Maeland AD, Bjordal M, Jensen MF, Hansen A, Flaat M, Weissenbach J, Lehrach H, Wincker P et al: Hox cluster disintegration with persistent anteroposterior order of expression in Oikopleura dioica. Nature 2004, 431(7004):67–71.

25. Serano JM, Martin A, Liubicich DM, Jarvis E, Bruce HS, La K, Browne WE, Grimwood J, Patel NH: Comprehensive analysis of Hox gene expression in the amphipod crustacean Parhyale hawaiensis. Developmental biology 2016, 409(1):297–309.

26. Irvine SQ, Martindale MQ: Expression Patterns of Anterior Hox Genes in the Polychaete Chaetopterus: Correlation with Morphological Boundaries. Developmental biology 2000, 217(2):333–351.

27. Lowe CJ, Wray GA: Radical alterations in the roles of homeobox genes during echinoderm evolution. Nature 1997, 389(6652):718–721.

28. Lee PN, Callaerts P, de Couet HG, Martindale MQ: Cephalopod Hox genes and the origin of morphological novelties. Nature 2003, 424(6952):1061–1065.

29. Godwin AR, Capecchi MR: Hoxc13 mutant mice lack external hair. Genes & development 1998, 12(1):11–20.

30. Aronowicz J, Lowe CJ: Hox gene expression in the hemichordate *Saccoglossus kowalevskii* and the evolution of deuterostome nervous systems. Integ Comp Biol 2006, 46(6):890–901.

31. Hejnol A, Martindale MQ: Coordinated spatial and temporal expression of Hox genes during embryogenesis in the acoel Convolutriloba longifissura. BMC biology 2009, 7:65.

32. Wada H, Garcia-Fernandez J, Holland PW: Colinear and segmental expression of amphioxus Hox genes. Developmental biology 1999, 213(1):131–141.

33. Chauvet S, Merabet S, Bilder D, Scott MP, Pradel J, Graba Y: Distinct hox protein sequences determine specificity in different tissues. Proceedings of the National Academy of Sciences of the United States of America 2000, 97(8):4064–4069.

34. Woltering JM, Duboule D: Tetrapod axial evolution and developmental constraints; Empirical underpinning by a mouse model. Mech Dev 2015, 138 Pt 2:64–72.

35. Zakany J, Duboule D: The role of Hox genes during vertebrate limb development. Curr Opin Genet Dev 2007, 17(4):359–366.

36. Wasik BR, Rose DJ, Moczek AP: Beetle horns are regulated by the Hox gene, Sex combs reduced, in a species- and sex-specific manner. Evolution & development 2010, 12(4):353–362.

37. Barucca M, Canapa A, Biscotti MA: An Overview of Hox Genes in Lophotrochozoa: Evolution and Functionality. J Dev Biol 2016, 4:12.

38. Dunn CW, Giribet G, Edgecombe GD, Hejnol A: Animal phylogeny and its evolutionary implications. Ann Rev Ecol Evol Syst 2014, 45:371–395.

39. Hejnol A: A Twist in Time-The Evolution of Spiral Cleavage in the Light of Animal Phylogeny. Integr Comp Biol 2010, 50(5):695–706.

40. Kocot KM: On 20 years of Lophotrochozoa. Org Divers Evol 2016:1–15.

41. Laumer CE, Bekkouche N, Kerbl A, Goetz F, Neves RC, Sørensen MV, Kristensen RM, Hejnol A, Dunn CW, Giribet G et al: Spiralian Phylogeny Informs the Evolution of Microscopic Lineages. Current biology: CB 2015, 25(15):2000–2006.

42. Struck TH, Wey-Fabrizius AR, Golombek A, Hering L, Weigert A, Bleidorn C, Klebow S, Iakovenko N, Hausdorf B, Petersen M et al: Platyzoan paraphyly based on phylogenomic data supports a noncoelomate ancestry of spiralia. Molecular biology and evolution 2014, 31(7):1833–1849.

43. Flot JF, Hespeels B, Li X, Noel B, Arkhipova I, Danchin EG, Hejnol A, Henrissat B, Koszul R, Aury JM et al: Genomic evidence for ameiotic evolution in the bdelloid rotifer Adineta vaga. Nature 2013, 500(7463):453–457.

44. Currie KW, Brown DD, Zhu S, Xu C, Voisin V, Bader GD, Pearson BJ: HOX gene complement and expression in the planarian Schmidtea mediterranea. EvoDevo 2016, 7:7.

45. Wasik K, Gurtowski J, Zhou X, Ramos OM, Delas MJ, Battistoni G, El Demerdash O, Falciatori I, Vizoso DB, Smith AD et al: Genome and transcriptome of the regeneration-competent flatworm, *Macrostomum lignano*. Proceedings of the National Academy of Sciences of the United States of America 2015, 112(40):12462–12467.

46. Fröbius AC, Matus DQ, Seaver EC: Genomic organization and expression demonstrate spatial and temporal Hox gene colinearity in the lophotrochozoan *Capitella* sp. I. PloS one 2008, 3(12):e4004.

47. Zhang G, Fang X, Guo X, Li L, Luo R, Xu F, Yang P, Zhang L, Wang X, Qi H et al: The oyster genome reveals stress adaptation and complexity of shell formation. Nature 2012, 490(7418):49–54.

48. Fritsch M, Wollesen T, de Oliveira AL, Wanninger A: Unexpected co-linearity of Hox gene expression in an aculiferan mollusk. BMC Evol Biol 2015, 15:151.

49. Fritsch M, Wollesen T, Wanninger A: Hox and ParaHox gene expression in early body plan patterning of polyplacophoran mollusks. Journal of experimental zoology Part B, Molecular and developmental evolution 2016, 326(2):89–104.

50. Hiebert LS, Maslakova SA: Expression of *Hox, Cdx*, and *Six3/6* genes in the hoplonemertean *Pantinonemertes californiensis* offers insight into the evolution of maximally indirect development in the phylum Nemertea. EvoDevo 2015, 6:26.

51. Hiebert LS, Maslakova SA: *Hox* genes pattern the anterior-posterior axis of the juvenile but not the larva in a maximally indirect developing invertebrate, *Micrura alaskensis* (Nemertea). BMC biology 2015, 13:23.

52. Hinman VF, O’Brien EK, Richards GS, Degnan BM: Expression of anterior Hox genes during larval development of the gastropod *Haliotis asinina*. Evolution & development 2003, 5(5):508–521.

53. Irvine SQ, Martindale MQ: Comparative analysis of Hox gene expression in the polychaete Chaetopterus: Implications for the evolution of body plan regionalization. Am Zool 2001, 41(3):640–651.

54. Kourakis MJ, Martindale MQ: Hox gene duplication and deployment in the annelid leech Helobdella. Evolution & development 2001, 3(3):145–153.

55. Kourakis MJ, Master VA, Lokhorst DK, Nardelli-Haefliger D, Wedeen CJ, Martindale MQ, Shankland M: Conserved anterior boundaries of Hox gene expression in the central nervous system of the leech *Helobdella*. Developmental biology 1997, 190(2):284–300.

56. Kourakis MJ, Master VA, Lokhorst DK, Nardelli-Haefliger D, Wedeen CJ, Martindale MQ, Shankland M: Evolutionary conservation of Hox gene expression in the CNS of the leech *Helobdella*. Developmental biology 1997, 186(2):B98–B98.

57. Samadi L, Steiner G: Involvement of Hox genes in shell morphogenesis in the encapsulated development of a top shell gastropod (*Gibbula varia* L.). Development genes and evolution 2009, 219(9–10):523–530.

58. Samadi L, Steiner G: Expression of Hox genes during the larval development of the snail, *Gibbula varia* (L.)-further evidence of non-colinearity in molluscs. Development genes and evolution 2010, 220(5–6):161–172.

59. Duboule D: Temporal colinearity and the phylotypic progression: a basis for the stability of a vertebrate Bauplan and the evolution of morphologies through heterochrony. Dev Suppl 1994:135–142.

60. Ferrier DEK, Minguillon C: Evolution of the Hox/ParaHox gene clusters. The International journal of developmental biology 2003, 47(7–8):605–611.

61. Monteiro AS, Ferrier DEK: Hox genes are not always Colinear. International journal of biological sciences 2006, 2(3):95–103.

62. Rudwick MJS: Living and fossil brachiopods: Hutchinson; 1970.

63. Brusca RC, Moore W, Shuster SM: Invertebrates. Sunderland, MA: Sinauer Associates, Inc.; 2016.

64. Luo YJ, Takeuchi T, Koyanagi R, Yamada L, Kanda M, Khalturina M, Fujie M, Yamasaki SI, Endo K, Satoh N: The *Lingula* genome provides insights into brachiopod evolution and the origin of phosphate biomineralization. Nat Commun 2015, 6:8301.

65. Freeman G: A developmental basis for the Cambrian radiation. Zoolog Sci 2007, 24(2):113–122.

66. Vellutini B, Hejnol A: Expression of segment polarity genes in brachiopods supports a nonsegmental ancestral role of engrailed for bilaterians. In. Scientific Reports; 2016: 32387.

67. Balavoine G, de Rosa R, Adoutte A: Hox clusters and bilaterian phylogeny. Molecular phylogenetics and evolution 2002, 24(3):366–373.

68. Halanych KM, Passamaneck Y: A Brief Review of Metazoan Phylogeny and Future Prospects in Hox-Research. Amer Zool 2001, 41:629–639.

69. Passamaneck YJ, Halanych KM: Evidence from Hox genes that bryozoans are lophotrochozoans. Evolution & development 2004, 6(4):275–281.

70. Ferrier DEK, Holland PWH: *Ciona intestinalis* ParaHox genes: evolution of Hox/ParaHox cluster integrity, developmental mode, and temporal colinearity. Molecular phylogenetics and evolution 2002, 24(3):412–417.

71. Patel NH: Evolutionary biology: time, space and genomes. Nature 2004, 431(7004):28–29.

72. Tümpel S, Wiedemann LM, Krumlauf R: Hox genes and segmentation of the vertebrate hindbrain. Curr Top Dev Biol 2009, 88:103–137.

73. Sharpe J, Nonchev S, Gould A, Whiting J, Krumlauf R: Selectivity, sharing and competitive interactions in the regulation of Hoxb genes. Embo J 1998, 17(6):1788–1798.

74. Kulakova M, Bakalenko N, Novikova E, Cook CE, Eliseeva E, Steinmetz PRH, Kostyuchenko RP, Dondua A, Arendt D, Akam M et al: Hox gene expression in larval development of the polychaetes *Nereis virens* and *Platynereis dumerilii* (Annelida, Lophotrochozoa). Development genes and evolution 2007, 217(1):39–54.

75. Spitz F, Gonzalez F, Duboule D: A global control region defines a chromosomal regulatory landscape containing the HoxD cluster. Cell 2003, 113(3):405–417.

76. Nielsen C: The development of the brachiopod *Crania (Neocrania) anomala* (O. F. Müller) and its phylogenetic significance. Acta Zoologica 1991, 72:7–28.

77. Freeman G: Metamorphosis in the Brachiopod *Terebratalia*: Evidence for a Role of Calcium Channel Function and the Dissociation of Shell Formation from Settlement. Biol Bull 1993, 184:15–24.

78. Steinmetz PRH, Kostyuchenko RP, Fischer A, Arendt D: The segmental pattern of otx, gbx, and Hox genes in the annelid *Platynereis dumerilii*. Evolution & development 2011, 13(1):72–79.

79. Negre B, Ruiz A: HOM-C evolution in Drosophila: is there a need for Hox gene clustering? Trends Genet 2007, 23(2):55–59.

80. Acemel RD, Tena JJ, Irastorza-Azcarate I, Marletaz F, Gómez-Marín C, de la Calle-Mustienes E, Bertrand S, Diaz SG, Aldea D, Aury JM et al: A single three-dimensional chromatin compartment in amphioxus indicates a stepwise evolution of vertebrate Hox bimodal regulation. Nat Genet 2016, 48(3):336–341.

81. Budd GE, Jensen S: A critical reappraisal of the fossil record of the bilaterian phyla. Biol Rev Camb Philos Soc 2000, 75(2):253–295.

82. Gustus RM, Cloney RA: Ultrastructural Similarities Between Setae of Brachiopods and Polychaetes1. Acta Zoologica 1972, 53(2):229–233.

83. Lüter C: Ultrastructure of Larval and Adult Setae of Brachiopoda. Zool Anz 2000, 239:75–90.

84. Orrhage L: Light and electron microscope studies of some brachiopod and pogonophoran setae. Z Morph Tiere 1973, 74(4):253–270.

85. Halanych KM, Bacheller JD, Aguinaldo AM, Liva SM, Hillis DM, Lake JA: Evidence from 18S ribosomal DNA that the lophophorates are protostome animals. Science 1995, 267(5204):1641–1643.

86. Lüter C, Bartolomaeus T: The phylogenetic position of Brachiopoda - a comparison of morphological and molecular data. Zool Scripta 1997, 26:245–253.

87. Fischer A: Mesoderm formation and muscle development of *Platynereis dumerilii* (Nereididae, Annelida). Dissertation. Berlin: Freie Universität Berlin; 2010.

88. Layden MJ, Meyer NP, Pang K, Seaver EC, Martindale MQ: Expression and phylogenetic analysis of the *zic* gene family in the evolution and development of metazoans. EvoDevo 2010, 1(1):12.

89. Smith MR: Ontogeny, morphology and taxonomy of the soft-bodied cambrian ‘mollusc’ *Wiwaxia*. Palaeontology 2014, 57:215–229.

90. Freeman G: Regional specification during embryogenesis in the articulate brachiopod *Terebratalia*. Developmental biology 1993, 160(1):196–213.

91. Freeman G: Regional Specification during Embryogenesis in the Craniiform Brachiopod *Crania anomala*. Developmental biology 2000, 227:219–238.

92. Reed C: Phylum Brachiopoda. In: Strathmann MF, editor. Reproduction and Development of the Marine Inverte-brates of the Northern Pacific Coast. Seattle. University of Washington Press 1987:pp 486–493.

93. Martin M: Cutadapt removes adapter sequences from high-throughput sequencing reads. *2011* 2011, 17(1).

94. Kajitani R, Toshimoto K, Noguchi H, Toyoda A, Ogura Y, Okuno M, Yabana M, Harada M, Nagayasu E, Maruyama H et al: Efficient *de novo* assembly of highly heterozygous genomes from whole-genome shotgun short reads. Genome Res 2014, 24(8):1384–1395.

95. Zimin AV, Marcais G, Puiu D, Roberts M, Salzberg SL, Yorke JA: The MaSuRCA genome assembler. Bioinformatics 2013, 29(21):2669–2677.

96. Bolger AM, Lohse M, Usadel B: Trimmomatic: a flexible trimmer for Illumina sequence data. Bioinformatics 2014, 30(15):2114–2120.

97. Grabherr MG, Haas BJ, Yassour M, Levin JZ, Thompson DA, Amit I, Adiconis X, Fan L, Raychowdhury R, Zeng Q: Full-length transcriptome assembly from RNA-Seq data without a reference genome. Nature biotechnology 2011, 29(7):644–652.

98. Katoh K, Standley DM: MAFFT multiple sequence alignment software version 7: improvements in performance and usability. Molecular biology and evolution 2013, 30(4):772–780.

99. Darriba D, Taboada GL, Doallo R, Posada D: ProtTest 3: fast selection of best-fit models of protein evolution. Bioinformatics 2011, 27(8):1164–1165.

100. Stamatakis A: RAxML version 8: a tool for phylogenetic analysis and post-analysis of large phylogenies. Bioinformatics 2014, 30(9):1312–1313.

101. Hejnol A, Martindale MQ: Acoel development indicates the independent evolution of the bilaterian mouth and anus. Nature 2008, 456(7220):382–386.

102. Santagata S, Resh C, Hejnol A, Martindale MQ, Passamaneck YJ: Development of the larval anterior neurogenic domains of *Terebratalia transversa* (Brachiopoda) provides insights into the diversification of larval apical organs and the spiralian nervous system. EvoDevo 2012, 3.

103. Grande C, Martín-Durán JM, Kenny NJ, Truchado-García M, Hejnol A: Evolution, divergence and loss of the Nodal signalling pathway: new data and a synthesis across the Bilateria. The International journal of developmental biology 2014, 58(6–8):521–532.

